# *Vibrio cholerae* El Tor strains linked to global cholera are homogeneous by pulsed-field gel electrophoresis

**DOI:** 10.1101/2021.11.22.469504

**Authors:** Fatema-Tuz Johura, Sahitya Ranjan Biswas, Shah M Rashed, Mohammad Tarequl Islam, Saiful Islam, Marzia Sultana, Anwar Huq, Nicholas R Thomson, Rita R. Colwell, Haruo Watanabe, Munirul Alam

## Abstract

*Vibrio cholerae* O1 El Tor, causative agent of the ongoing seventh cholera pandemic, is native to the aquatic environment of the Ganges Delta, Bay of Bengal (GDBB). Recent studies traced pandemic strains to the GDBB and proposed global spread of cholera had occurred via intercontinental transmission. In the research presented here, *Not*I-digested genomic DNA extracted from *V. cholerae* O1 clinical and environmental strains isolated in Bangladesh during 2004 – 2014 was analyzed by pulsed-field gel electrophoresis (PFGE). Results of cluster analysis showed 94.67% of the *V. cholerae* isolates belonged to clade A and included the majority of clinical isolates of spatio-temporal origin and representing different cholera endemic foci. The rest of the strains were estuarine, all environmental isolates from Mathbaria, Bangladesh, and occurred as singletons, clustered in clades B and C, or in the small clades D and E. Cluster analysis of the Bangladeshi strains and including 157 El Tor strains from thirteen countries in Asia, Africa, and the Americas revealed 85% of the total set of isolates belonged to clade A, indicating all were related, yet did not form an homogeneous cluster. Overall, 15% of the global strains comprised multiple small clades or segregated as singletons. Three sub-clades could be discerned within the major clade A, reflecting distinct lineages of *V. cholerae* El Tor associated with cholera in Asia, Africa, and the Americas. The presence in Asia and the Americas of non-pandemic *V. cholerae* El Tor populations differing by PFGE and from strains associated with cholera globally suggests different ecotypes are resident in distant geographies.

**Author Summary:** Cholera is a major health threat, especially in the Ganges Delta, Bay of Bengal (GDBB). *Vibrio cholerae*, causative agent of cholera, is native to the GDBB aquatic environment. Recent genomic studies suggest GDBB is the cholera hotspot where the disease spreads globally via human activity. Pulsed-field gel electrophoresis (PFGE) of *Not*I-digested genomic DNA from *V. cholerae* El Tor endemic cholera strains was done, including Bangladesh aquatic environment and clinical strains from distant geographical regions representing three cholera-prone continents. Results showed the majority of pandemic strains belonged to a major cluster, suggesting clonal relatedness. Ecotypes were detected, indicating geographically specific lineages. It is concluded that epidemic strains in Bangladesh and thirteen countries of Asia, Africa, and the Americas are geographically adapted, with independent evolution of the bacterium in respective geographical regions.

## Introduction

Cholera, with seven pandemics reported to date, represents a significant chapter in human history and infectious disease. The acute form of diarrhea caused by *Vibrio cholerae* is related to production of a toxin that triggers the characteristic water loss and severe dehydration of cholera [1]. Cholera remains a threat in many countries, notably where access to safe drinking water is limited. Bangladesh is a developing country where cholera is endemic, with two annual peaks in some regions of the country [2]. The estimated global burden of cholera is 2.86 million cases and 95,000 deaths. In Bangladesh alone, approximately 100,000 million new cases are diagnosed each year, resulting in 4,500 deaths [3]. *V. cholerae* strains are widely distributed globally in many coastal, estuarine, and brackish water ecosystems as free-living bacterial cells or associated with zooplankton, namely copepods [4-6]. Brackish waters in coastal areas support bacterial populations, with environmental stimuli favorable for bacterial growth prompting cholera outbreaks [5].

*V. cholerae* O1 is classified into two biotypes, classical and El Tor, based on genetic differences. The seventh and ongoing pandemic is attributed to the El Tor biotype of *V. cholerae* O1 [8]. Beginning in 1961 in Indonesia, the seventh pandemic of cholera included Africa in 1970, Latin America in 1991, and more recently Haiti and Yemen [9-12]. After the initial cases occurred, most of these regions continued to suffer episodes of cholera. There is debate whether the recurrent outbreaks of cholera in Africa and the Americas are caused by distinct intercontinental introduction or resurgence of indigenous clones. A few recent investigations investigated the genetic homogeneity of the 7^th^ pandemic strains and connected both origin and recurrent transmission to a single source, the Bay of Bengal [13]. However, the co-existence of several local lineages, along with the pandemic clones and their regional evolution, has painted a very complex picture of bacterial population dynamics, especially in and out of endemic settings [14]. Thus, monitoring pandemic and local strains became a priority for some investigators, as a means of developing an effective public health model and strategy for controlling the current pandemic and preventing future pandemics of cholera.

Whole genome typing methods, e.g. multi-locus sequence typing (MLST), multi-locus variant analysis (MLVA), ribotyping, random amplification of polymorphic DNA (RAPD), and pulsed field gel electrophoresis (PFGE), have been employed to differentiate isolates and monitor transmission routes [15-18]. PFGE is a DNA fingerprinting method that can discriminate bacterial strains. Before introduction of whole genome sequencing (WGS), epidemiological studies of cholera relied on PFGE [18]. An earlier study highlighted the value of PFGE in revealing clonality among isolates from two well-defined cholera outbreaks in Malaysia [19]. Intrinsic limitations include restriction digestion being skewed by mobile elements, hence restricted value for phylogeny. Although PFGE does not provide as high resolution as WGS, its stability and reproducibility allow rudimentary, yet comprehensive, analysis of ancestry [18].

In the study reported here, the objective was to understand both regional diversity and global distribution of *V. cholerae* El Tor representing seventh pandemic lineage. Therefore, strains in Bangladesh isolated over a decade were compared with strains from thirteen countries across Asia, Africa, and the Americas. Both environmental and clinical isolates were included since the local environment can influence strains, persisting and becoming epidemic, as well as providing access to autochthonous strains of *V. cholerae* El Tor.

## Results

The *Not*I restriction enzyme digested genomic DNA of the test strains into 20 to 23 fragments and the molecular sizes of the DNA fragments ranged from 20 to 350 kb. Digested genomic DNAs of different spatiotemporal origin and resulting different biotype categorizations were subjected to PFGE. The resulting band patterns were analyzed by Dice similarity coefficient and UPGMA clustering methods to determine genetic and ancestral relatedness.

### Local diversity and distribution of the Bangladesh isolates

In a dendogram obtained by UPGMA analysis of DNA band patterns, the Bangladesh isolates comprised five different clades, A, B, C, D, and E (Figs 1 and 2). Of the 169 isolates, 160 clustered in clade A, suggesting a single lineage. A single clinical isolate, collected in 1991 from Matlab, clustered with clade A, a clade of predominantly clinical isolates from endemic sites, including estuary villages in Bangladesh. While a few environmental isolates were found to cluster in clade A, other environmental isolates collected in 2012 from Mathbaria exhibited different PFGE patterns and did not join clade A (S1 Fig). Isolates with different pulsotypes included three singletons and a small clade [B(1); C(1); D(6); E(1)] (Fig 1). Singletons in clades B, C, and E were isolates from the aquatic environment. It should be noted that clade D included isolates of both clinical and environmental origin. All clade D isolates possessed *rstR* classical biotype, a characteristic limited to this clade (S1 Table). Most of the Chhatak isolates (27 of 31) had the same pulsotype closely related to *V. cholerae* isolates from Dhaka and Mathbaria (S1 Fig).

**Fig 1.**
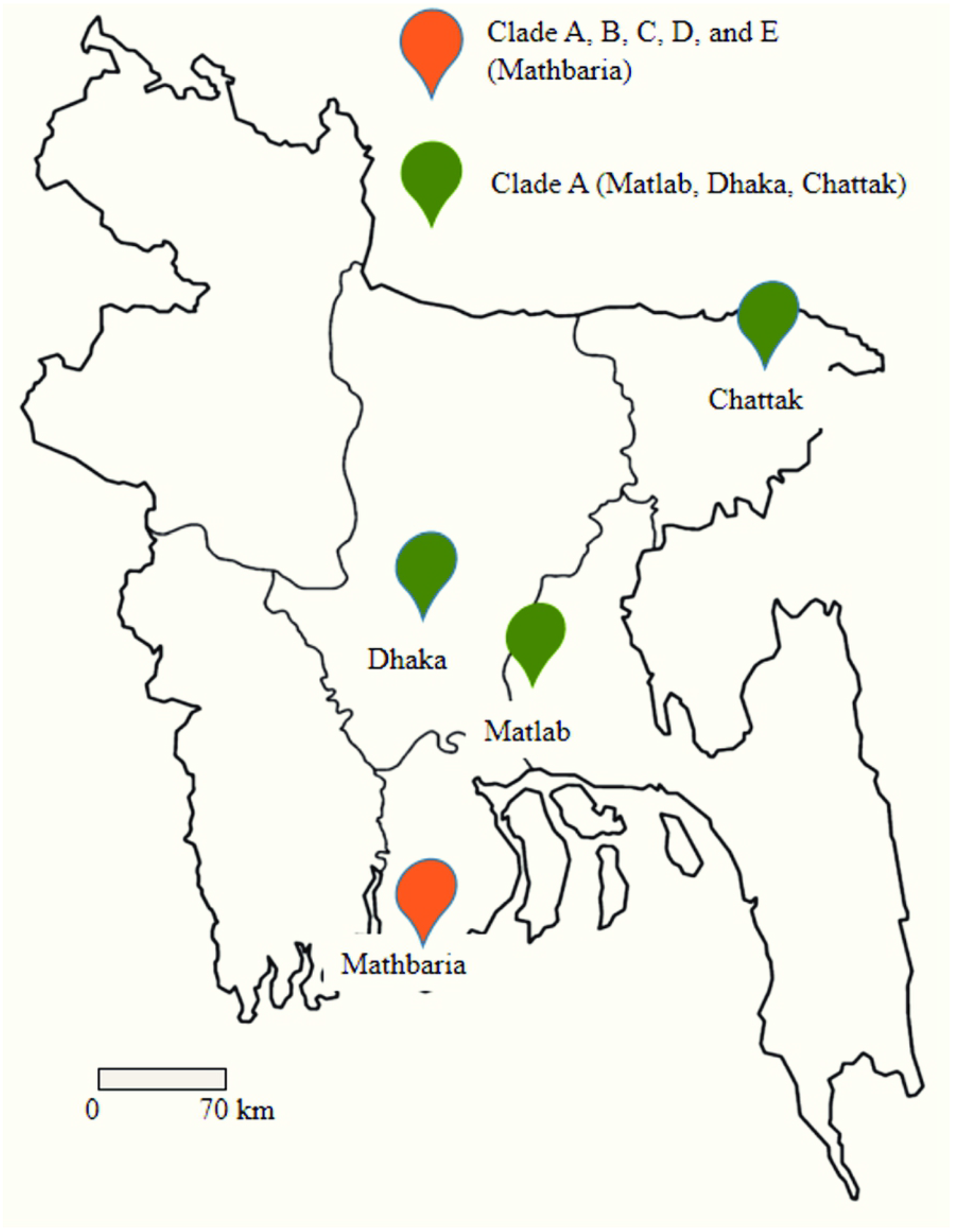
Clonal diversity and geographical distribution of isolates from Bangladesh. Isolates belonging to clade A were found in all four areas. Clade A contained both clinical and environmental isolates. In addition to clade A, strains of other clades were found, but only in Mathbaria.

**Fig 2.**
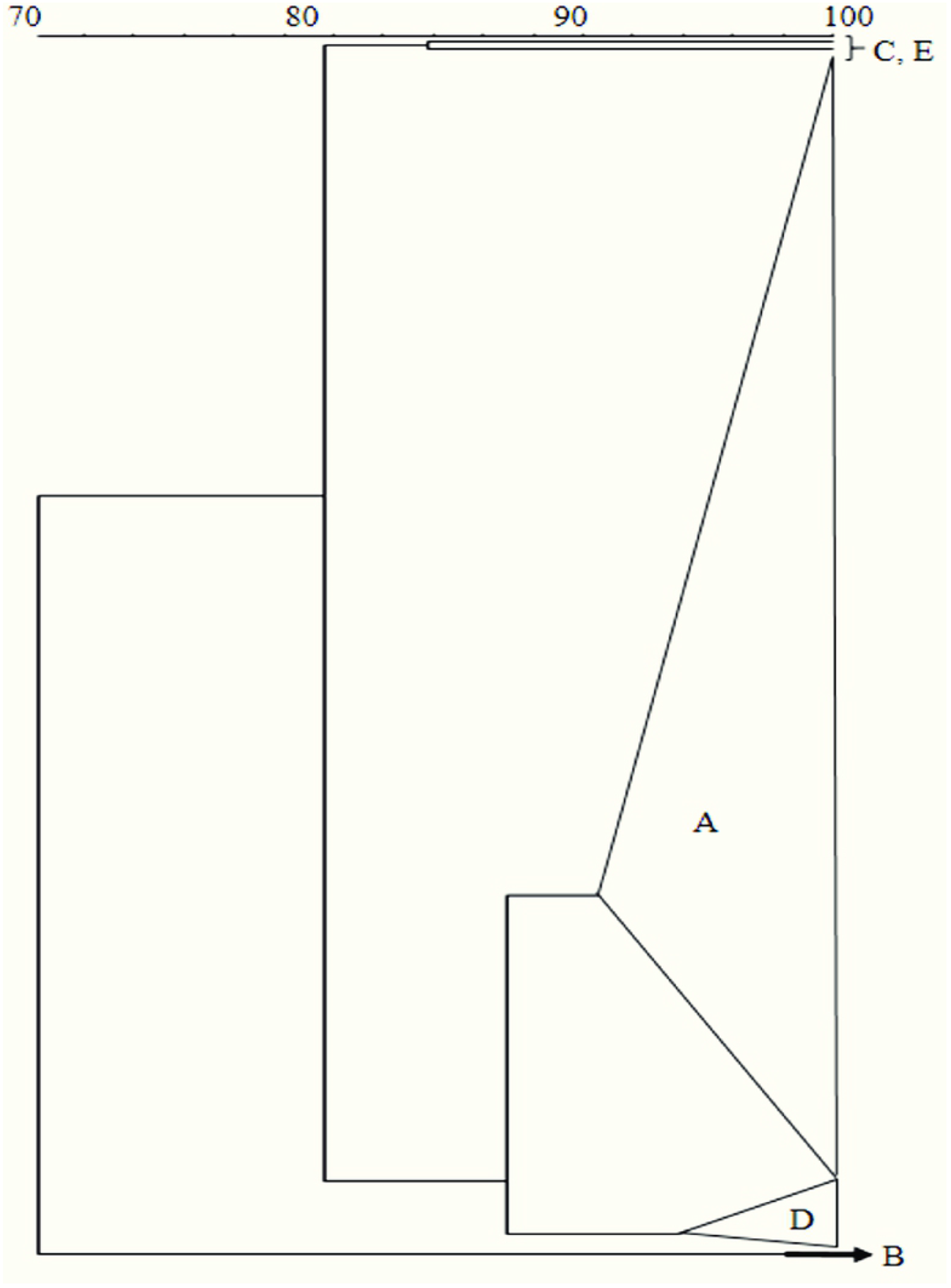
PFGE analysis of the isolates collected from Bangladesh. A total of 169 strains were analyzed which resulted in 5 clades. Clade A contained 94.67% of the isolates. The rest 9 isolates which were only found in Mathbaria formed the other 4 groups (B, C, D and, E).

### Global distribution of clones

When the PFGE banding patterns of *V. cholerae* O1 isolates from Bangladesh were compared with those of 157 isolates collected from thirteen countries across Asia, Africa, and Latin America, the isolates could be differentiated into 16 clades; A through P (Fig 3). Clade A isolates from Bangladesh clustered with 133 of the 293 strains, comprising a majority of isolates from 14 countries and three continents (Figs 3 and 4). Hence, a majority of the *V. cholerae* El Tor isolates from different geographical regions revealed a similar PFGE pattern and fell into a major clade, with country-specific sub-clustering, i.e., subclades within the major clade A. The sub clades of A included *V. cholerae* El Tor strains from Vietnam (n=15), Zambia (n=9), Haiti (n=3), India (n=2), Pakistan (n=3), Sri Lanka (n=1), and most of the strains from Zimbabwe (8 of 12) and Nepal (27 of 39), Fig 4. Three subclades within clade A reflected a broader spatial distinction and 202 of 273 (74%) Asian and African isolates comprising subclade Ia and 60 of 273 (22%) in subclade Ib. Latin American isolates (16 of 21) in clade A comprised subclade II. Interestingly, the Latin American isolates were predominantly *V. cholerae* prototype ET (S1 Table). A few isolates from Bangladesh, Thailand and one from Zimbabwe fell into sub-clade II (Fig 4). Subclade II *V. cholerae* El Tor strains from Mexico, isolated during 1992 to 1999, were located in a branch, separating them from isolates collected between 2004 and 2008. Three *V. cholerae* El Tor strains isolated during the 2010 Haitian cholera outbreak comprised subclade Ia, with Asian and African strains. Two *V. cholerae* O1 EL Tor strains isolated in Bangladesh during 2011 joined with *V. cholerae* ET strains from Peru, Brazil, and Mexico, based on PFGE, Figs 3 and 4.

**Fig 3.**
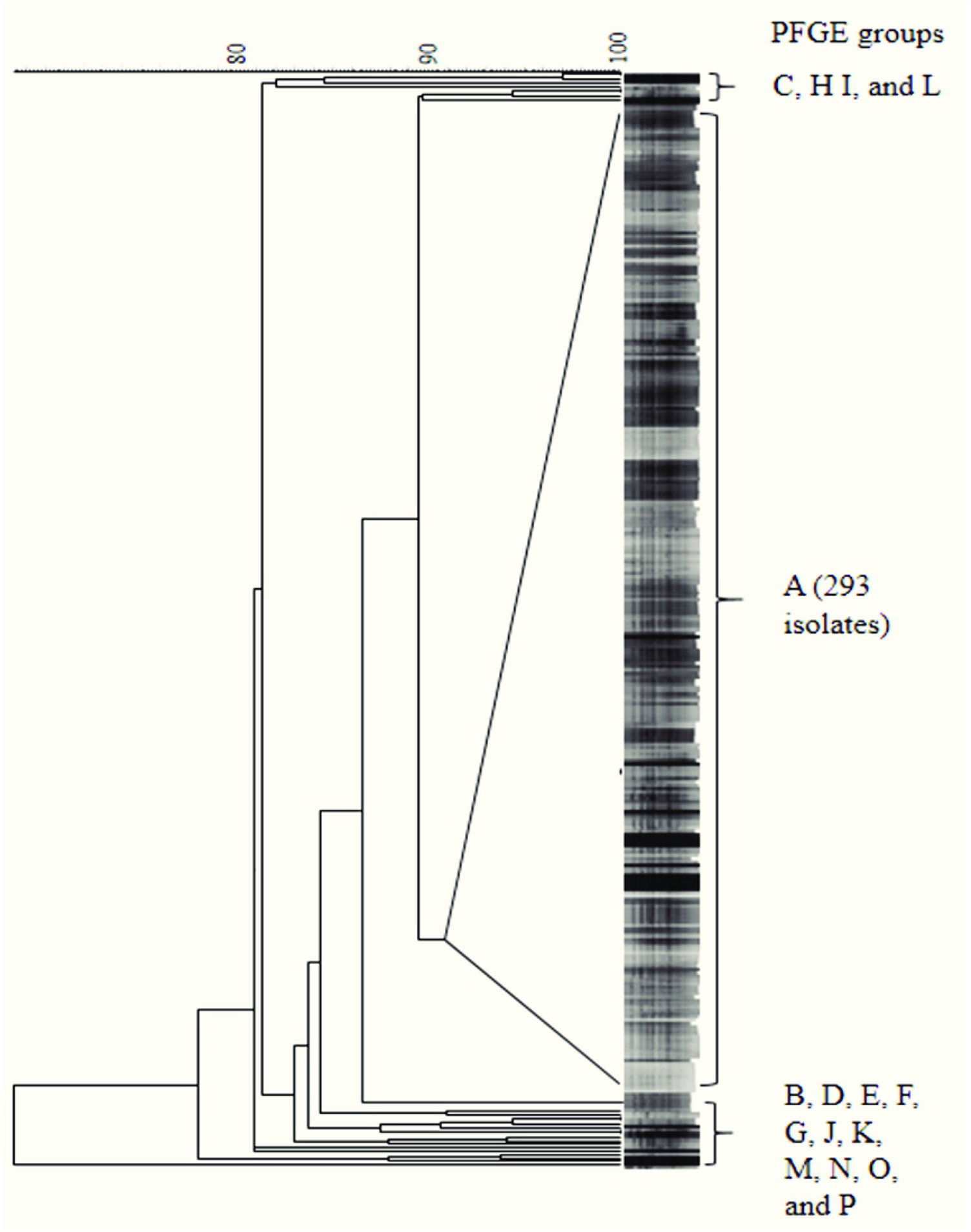
Comparison of band patterns between isolates from Bangladesh and other countries. The isolates from Bangladesh were compared with 157 additional isolates collected from 13 other countries. The resulting phylogenetic tree represents 16 groups. Clade A contained 89.88% of the total isolates including 160 isolates of Bangladesh.

**Fig 4.**
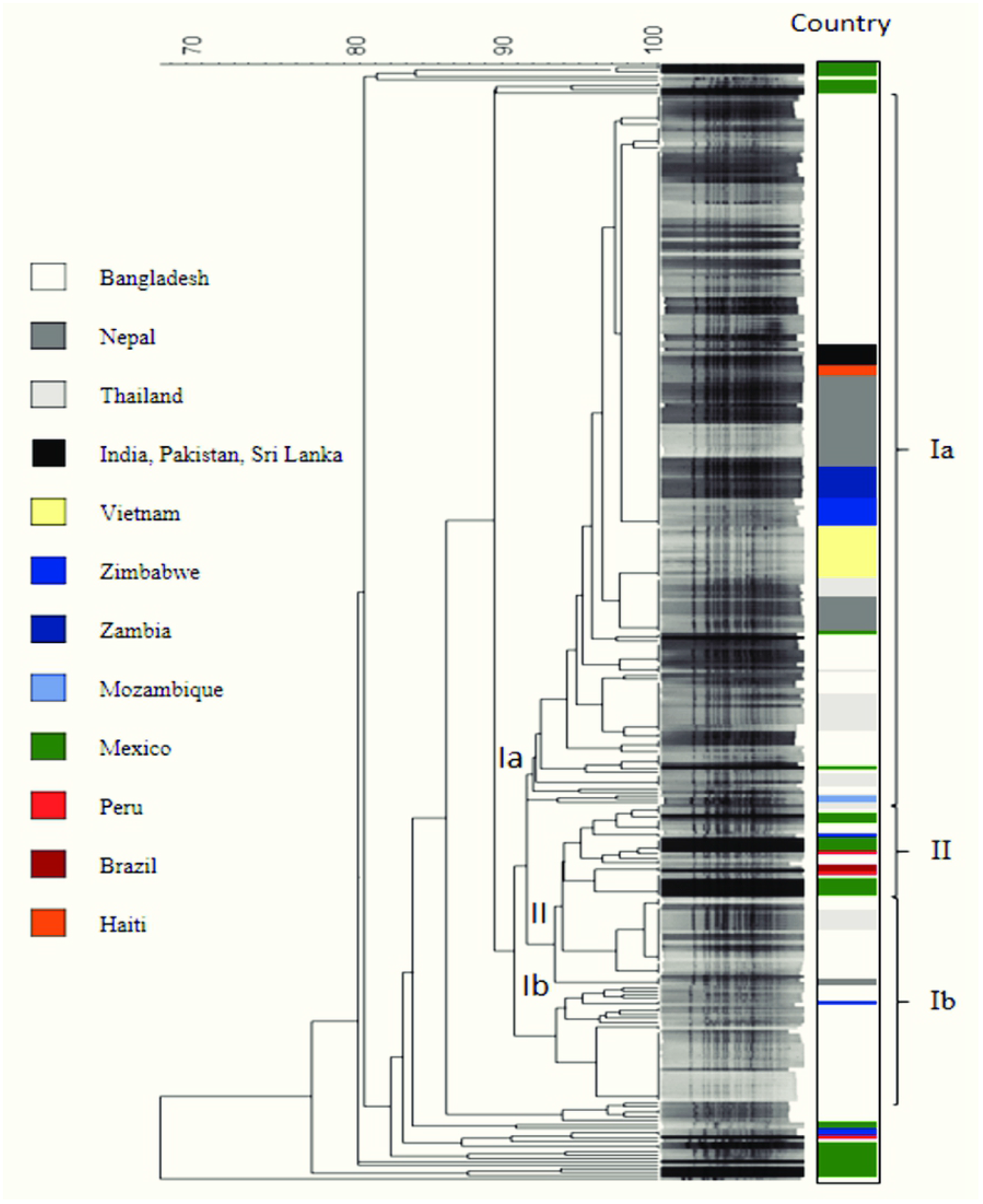
Global phylogeny of isolates based on PFGE pattern. Countries are represented by color. 160 isolates of Bangladesh formed clade A, with 133 strains isolated from other countries. Three subclades were observed in clade A: Subclade Ia and Ib contained strains mostly of African and Asian origin. Subclade II comprised predominantly Latin American strains. Country subclusters were also observed.

Aside from clade A, the clade E isolates from Bangladesh shared PFGE pattern with an ET isolate from Peru and two from Zimbabwe (Fig 5). As with the Bangladesh isolates, locally restricted diversity was observed for isolates from Mexico, reflected by 11 distinct groups in addition to clade A. Groups F-P comprised isolates from Mexico, notably those collected during 2000 to 2004 (Fig 4).

**Fig 5.**
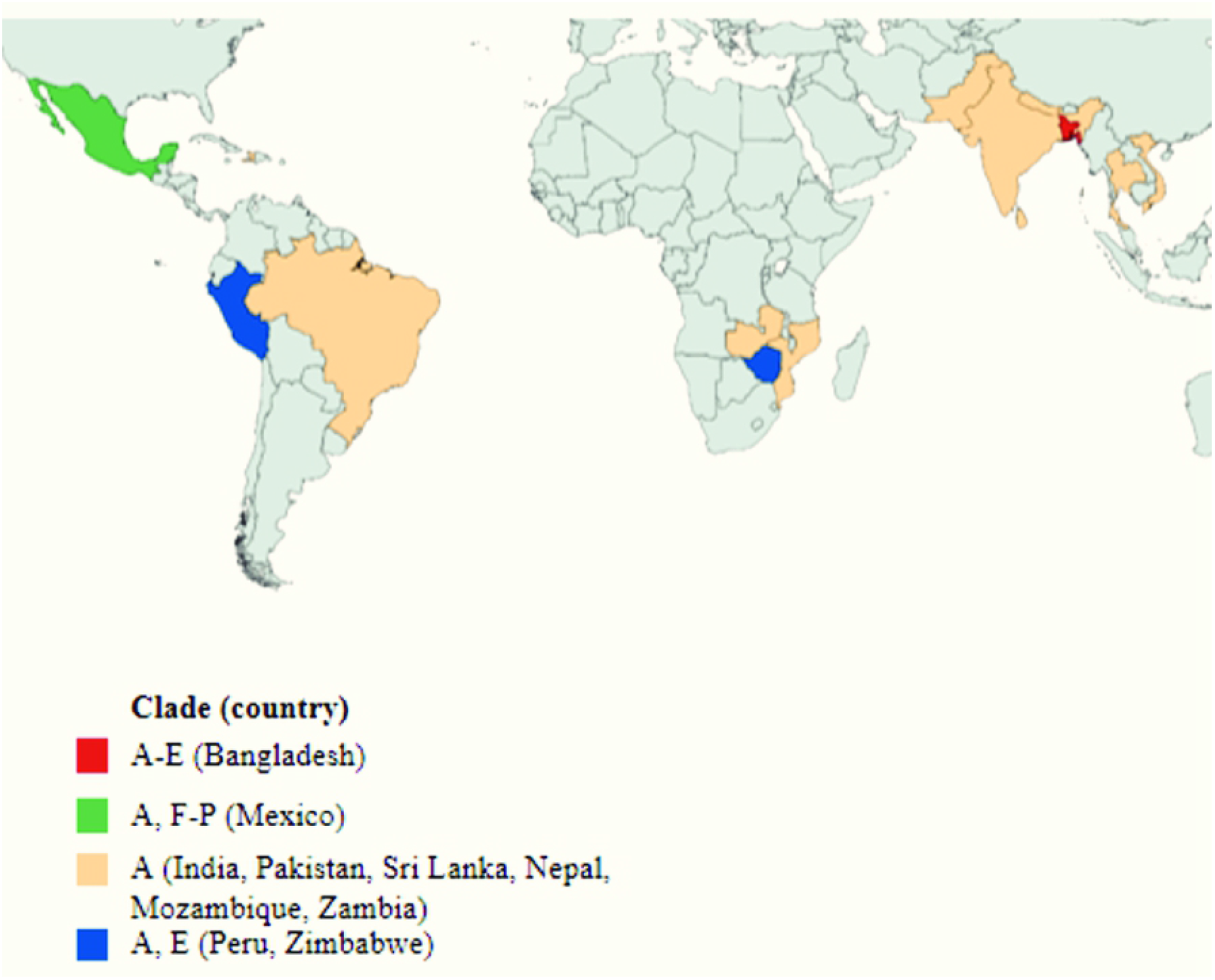
Global distribution of clones.

## Discussion

In many epidemiological studies, pulsed-field gel electrophoresis (PFGE) is used to discern source attributes of strains from different outbreaks. With the advent of whole genome sequencing and comparative genomics, the once gold standard PFGE is less appreciated as a DNA fingerprinting tool of epidemiological implication. In the study reported here, PFGE analysis of *V. cholerae* O1 El Tor isolates associated with endemic cholera in Bangladesh and thirteen countries of Asia, Africa and the America showed the strains to be related genetically, but not homogenous globally. A majority of the strains comprised a major clade, with divergence noted for non-pandemic and environmental isolates from the Bay of Bengal, Bangladesh and strains from the Gulf of Mexico. According to a recent WGS-based study of a restricted subset of *V. cholerae* clones, those strains were responsible for epidemic cholera worldwide [20-21]. In this study, PFGE was used to analyze clinical strains of *V. cholerae* from different geographical locations and with different genetic and phenotypic characteristics.

*V. cholerae* El Tor biotype has dominated clinically over the classical biotype since 1961, the latter having last been isolated in Bangladesh in 1992 [22-23]. Observed co-existence of the two biotypes for such a long time likely resulted in the hybrid characteristics of El Tor with classical biotype attributes, as observed in Bangladesh [24-25]. In this study, the majority of *V. cholerae* El Tor isolates from clinical and environment sources comprised a major clade, suggesting similarity of strains from both sources and associated with epidemics in Bangladesh. Some of the environmental strains did not fall into clade A, suggesting those to be genetically divergent pulsotypes present in a diverse population existing in the environment. Environmental *V. cholerae* ET in our study, with a few exceptions, were similar to clinical isolates in clade A. Previous epidemiological surveillance conducted in the Bay of Bengal estuary has shown pathogenic strains can be detected in aquatic habitats, either in the culturable or non-culturable state, depending on the season [26].

A major genome-based study postulated the pandemic *V. cholerae* strain originated in Bay of Bengal villages of Asia and transmitted world-wide in three different waves [13]. It was concluded that *V. cholerae* O1 ET has the ability to travel inter-continentally and adapt to its place of introduction by sharing niches with existing microflora in coastal and estuarine regions, including the Gulf coast of Mexico [27]. Notwithstanding the fact that outbreaks occurring after introduction can be attributed to *V. cholerae* and the pathogen can be introduced repeatedly or adapt locally, either is possible. Whole genome sequencing based studies linked epidemics in Africa and the Americas to multiple introduction events, rather than preexisting pathotypes [8-9]. The PFGE banding patterns observed for the majority of *V. cholerae* O1 included in this study support an intercontinental transmission hypothesis [28], but only in a global context. The observation of country-based subclades indicates an independent evolution of the pandemic pathogen. Genetic changes were reported among initially homogeneous *V. cholerae* O1 ET initiating the Haitian cholera epidemic in 2010 [28]. In this context, *V. cholerae* ET strains associated with the Haitian cholera in 2010 were observed to be closely related to isolates from other Southeast Asian countries [28].

While cholera had not been reported in the Americas for more than a century before 1991, the observed presence of *V. cholerae* classical biotype and diverse *V. cholerae* ET lineages in Mexico was uncharacteristic for a region outside of Asia or Africa, suggesting a capricious nature of the bacterium [29]. Most *V. cholerae* ET isolates in Mexico that diverged separately from the pandemic clones lacked CTX phage [30] and were not related to the non-toxigenic isolates from Thailand [31]. Previously, we had shown that CTX phage negative isolates dominated clinical cases in Mexico during 2001-2004 [30] and studies posited the isolates to be ancestors of the *V. cholerae* responsible for the sixth and seventh pandemics [29].

While the observed relatedness of PFGE patterns of *V. cholerae* El Tor associated with cholera epidemics in Asia, Africa, and the Americas supports the potential for global transmission of the pandemic pathogen [13], the divergence of strains and their region-specific signatures also support independent evolution of *V. cholerae* locally. Clearly the overall picture is complex and warrants regular monitoring to assist designing effective intervention models to counter future pandemics. In any case PFGE data, as presented in this study show this technology can be effective for analysis and source-tracking of cholera outbreaks.

## Materials and Methods

### Geographical profile of isolates

A total of 169 strains were collected between 1991 and 2014 from four endemic sites in Bangladesh: Mathbaria (n=99); Dhaka (n=38); Chhatak (n=31); and Matlab (n=1). Of these, 119 and 50 were of clinical and environmental origin, respectively. To investigate phylogenetic relationships, an additional 157 strains (150 clinical and 7 environmental) were collected from 13 countries across Asia, Africa, and Latin America [Nepal 39, Thailand 32, Vietnam 15, Pakistan 3, India 2, Sri Lanka 1, Zambia 9, Zimbabwe 12, Mozambique 2, Mexico 34, Brazil 2, Peru 3, and Haiti 3] (Table 1). All strains were confirmed as *V. cholerae* O1 El Tor biotype by culture, and serotype and biotype specific genotype. Detailed information for the isolates is provided in Tables 1 and 2, and S1 Table.

**Table 1.** Geographic source of *V. cholerae* El tor isolates included in this study.

| Country | No. of strains | Source |  | Serotype |  |
| --- | --- | --- | --- | --- | --- |
|  |  | Environmental | Clinical | Inaba | Ogawa |
| Bangladesh | 169 | 50 | 119 | 33 | 136 |
| Sri Lanka | 1 | - | 1 | - | 1 |
| Pakistan | 3 | - | 3 | 1 | 2 |
| India | 2 | - | 2 | 2 | - |
| Nepal | 39 | 6 | 33 | - | 39 |
| Vietnam | 15 | - | 15 | - | 15 |
| Thailand | 32 | 1 | 31 | 17 | 15 |
| Zambia | 9 | - | 9 | - | 9 |
| Zimbabwe | 12 | - | 12 | 4 | 8 |
| Mozambique | 2 | - | 2 | - | 2 |
| Mexico | 34 | - | 34 | 17 | 17 |
| Brazil | 2 | - | 2 | 2 | - |
| Haiti | 3 | - | 3 | - | 3 |
| Peru | 3 | - | 3 | - | 3 |
| n= 326 |  |  |  |  |  |

**Table 2.** *V. cholerae* genotypes based on *ctxB, rstR, and tcpA* genes.

| Countries | *ctxB* |  |  | *rstR* |  |  |  | *tcpA* |  |  |  |
| --- | --- | --- | --- | --- | --- | --- | --- | --- | --- | --- | --- |
|  | Classical | El Tor | Negative | Classical | El Tor | Both | Negative | Classical | El Tor | Both | Negative |
| Bangladesh | 167 | 2 | - | 7 | 162 | - | - | - | 169 | - | - |
| Nepal | 39 | - | - | - | 39 | - | - | - | 39 | - | - |
| Thailand | 27 | - | 5 | 9 | 18 | - | 5 | - | 32 | - | - |
| India | 2 | - | - | - | 2 | - | - | - | 2 | - | - |
| Pakistan | 3 | - | - | - | 3 | - | - | - | 3 | - | - |
| Sri Lanka | 1 | - | - | - | 1 | - | - | - | 1 | - | - |
| Vietnam | 15 | - | - | - | 15 | - | - | - | 15 | - | - |
| Zambia | 9 | - | - | - | 9 | - | - | - | 9 | - | - |
| Mozambique | 2 | - | - | 2 | - | - | - | - | 2 | - | - |
| Zimbabwe | 12 | - | - | - | 12 | - | - | - | 12 | - | - |
| Mexico | 5 | 16 | 13 | - | 16 | 5 | 13 | 1 | 26 | 4 | 3 |
| Brazil | - | 2 | - | - | 2 | - | - | - | 2 | - | - |
| Peru | - | 3 | - | - | 3 | - | - | - | 3 | - | - |
| Haiti | 3 | - | - | - | 3 | - | - | - | 3 | - | - |

### Pulsed-field gel electrophoresis (PFGE)

Whole agarose-embedded genomic DNA was prepared from each isolate. PFGE was carried out using a contour-clamped homogeneous electrical field (CHEF-DRII) apparatus (Bio-Rad), following procedures described previously [32]. Conditions for separation were as follows: 2 to 10s for 13 h, followed by 20 to 25 s for 6 h. An electrical field of 6 V/cm was applied at an included field angle of 120°. Genomic DNA of the test strains was digested by *Not*I restriction enzyme (Gibco-BRL, Gaithersburg, MD) and *Salmonella enterica* serovar Braenderup was digested using *Xba*I, with fragments employed as molecular size markers. Restriction fragments were separated in 1% pulsed-field-certified agarose in 0.5X TBE (Tris-borate-EDTA) buffer. Post-electrophoresis gel-treatment included gel staining and de-staining. DNA was visualized using a UV transilluminator and images were digitized using a one-dimensional gel documentation system (Bio-Rad).

### Image analysis

The fingerprint pattern in each gel was analyzed using a computer software package, Bionumeric (Applied Maths, Belgium). After background subtraction and gel normalization, the fingerprint patterns were typed according to banding similarity and dissimilarity, using the Dice similarity coefficient and unweighted-pair group method employing average linkage (UPGMA) clustering, as recommended by the manufacturer. The results were graphically represented as dendrograms.

### Institutional approval

All the experimental protocols were reviewed and approved by the Research Review Committee (RRC), and Ethics Review Committee(ERC) of the International Centre for Diarrhoeal Disease Research, Bangladesh (research grant: 1R01A139129-01 and protocol: PR-14017). All methods were conducted in accordance with the guidelines of the RRC and ERC.

## Acknowledgement

This research was partially supported by Japan Food Hygiene Association through National Institutes of Infectious Diseases (NIID), Tokyo, Japan, and the NIH (USA) research grant 1R01A139129-01 under collaborative agreements between the Johns Hopkins Bloomberg School of Public Health, icddr,b, and the University of Maryland. Authors gratefully acknowledge Richard Bradley Sack, Johns Hopkins Bloomberg School of Public Health, USA, and K-M Kam and Cindy KY Luey, Public Health Laboratory Center, Hong Kong, and the icddr,b hospital and laboratory staff for their support. icddr,b gratefully acknowledges the following donors who provide unrestricted support: Governments of Bangladesh, Canada, Sweden and the UK.

## Author Contributions

Conceptualization and study design: FTJ, MA

Data curation: FTJ, SRB

Formal analysis: FTJ, SRB

Investigation: FTJ, SMR, MTI, SI, MS

Methodology: FTJ, SMR, MTI, SI, MA

Supervision: MA

Writing – original draft: FTJ, SRB

Writing—review and editing: TA, AH, NRT, RRC, MA

All authors contributed to the article and approved the submitted version

## Supporting information

**S1 Fig. Cluster analysis of isolates from Bangladesh**. The area and year of isolation are shown in color codes.

**S1 Table. *Vibrio cholerae* El Tor strains included in this study with source and year of isolation**.

